# Spatially resolved, multimodal in vivo Perturb-seq using antibody-based cell hashing

**DOI:** 10.64898/2026.05.25.727765

**Authors:** Alexander A Nevue, George C Hartoularos, Cynthia De Valle, Kheerthivasan Ramachandran, Jerika J Barron, Maria Erendira Calleja Cervantes, Hannah Lee, Joseph Bowness, Lars Velten, Chiara Ricci-Tam, Maayan Levy, Alexander Dobin, Chun Jimmie Ye, Inna Averbukh, David Lara-Astiaso

**Author notes:** these authors contributed equally to the work (Alexander A Nevue, George C Hartoularos). these authors jointly supervised the work (Inna Averbukh, David Lara-Astiaso).

## Abstract

Large-scale perturbation screens have begun to map cell intrinsic gene-function relationships, yet how genes shape tissue architecture remains largely unexplored. To address this gap, we developed PerturbSpace, a novel approach that integrates CRISPR perturbations with spatially hashed single-cell multiomics. This approach enables the first high-throughput, spatially resolved Perturb-seq analysis across complex tissue architecture in vivo. Notably, PerturbSpace enables spatial transcriptome-wide perturbation readouts at organ scale and can be seamlessly integrated with orthogonal modalities. We combine PerturbSpace with surface proteomics and expressed lineage tracing barcodes to demonstrate multimodal compatibility. We use PerturbSpace to study the genetic determinants of tissue architecture. First, we map how 40 transcriptional regulators determine the size and lineage composition of colonies in the spleen during regenerative hematopoiesis. Second, we characterize immune-niche interactions in the liver by dissecting the extrinsic effects mediated by cytokine-secreting immune cells on their neighboring cells. Collectively, our work establishes PerturbSpace as a scalable and cost-effective approach for transcriptome-wide spatial profiling of whole cells while remaining compatible with the single-cell multiomics workflows that the field has already adopted at scale.

## Introduction

In vivo Perturb-seq is emerging as a powerful tool to map gene functions across cells in their true physiological context^1^. However, this workflow relies on single-cell profiling that requires dissociation, which inherently destroys tissue contexts, limiting the analysis to cell intrinsic effects. Losing spatial context hinders the investigation of how genes regulate cellular interactions and cooperativity to establish tissue organization and function. Furthermore, the absence of tissue context limits our ability to interrogate how gene functions are modulated by the environmental conditions found within diverse tissue microenvironments. Recent spatial innovations, including optical and sequencing-based approaches, have begun to bridge this gap by enabling the profiling of perturbed cells within intact tissues^2–10^. These methods have provided valuable links between gene function and spatial features such as cellular morphology, but they lack the scale and modularity needed to systematically dissect the roles of genes in tissue organization, particularly the non-cell-autonomous effects that sustain tissue structure.

To address these limitations, we developed PerturbSpace, an unbiased multimodal readout of perturbed single cells at tissue neighborhood resolution using a spatial hashing approach. PerturbSpace requires minimal infrastructure, utilizes commercial single cell workflows, and is compatible with multimodal readouts. We apply PerturbSpace to interrogate the genetic determinants of tissue function in two paradigms. First, we characterize how genetic perturbations influence the composition and size of splenic colonies during regenerative hematopoiesis. Second, we map the non-cell-autonomous effects mediated by cytokine-producing immune cells within their specific tissue niches in the liver.

## Results

### Development of PerturbSpace for spatially resolved multimodal in vivo Perturb-seq

Functional genomics in tissue biology demands ultra-high-throughput spatial profiling for two reasons. First, *in vivo* perturbed cells are sparse, so capturing them in sufficient numbers requires large tissue areas. Second, resolving how perturbations reshape tissue architecture requires high scale profiling of both perturbed cells and their unperturbed neighbors while retaining their spatial context. To meet this need, we developed PerturbSpace, a spatially resolved functional genomics method that pairs a novel tissue hashing strategy with high-throughput single-cell multiomics. PerturbSpace jointly captures whole transcriptomes, sgRNAs, and spatial coordinates at organ scale offering four key advantages over other existing spatial perturbation readouts: i) requires minimal instrumentation and hands-on time enabling organ-wide profiling ii) generates a single-cell RNA-seq dataset facilitating harmonization with existing atlases iii) enables enrichment of spatially tagged cells via FACS maximizing utility of the final dataset, and iv) can be extended to obtain orthogonal multimodal information including surface proteomics (e.g. CITE-seq^11^), epigenetic (e.g. ATAC-seq), TCR/BCR repertoire, and genetic perturbations (e.g. Perturb-seq^12–14^).

To achieve scalable spatial barcoding of tissues, we developed high-density microwell arrays featuring spatial indexes conjugated to a tissue labelling reagent consisting of *a*MHC-I and *a*CD45 antibodies. Because these surface markers are present on all nucleated mammalian cell types, this approach enables universal spatial hashing of tissue sections. Following spatial hashing, tissues are dissociated into single-cell suspensions, and the spatially hashed cells are isolated by FACS. Then, by using a single-cell sequencing workflow designed to recover both the transcriptomes and their original spatial coordinates, we can achieve a seamless reconstruction of tissue architecture at 80µm resolution (Fig. 1a, Extended Data Fig. 1a-c). We benchmarked this approach in the kidney and liver demonstrating that this method produces high-quality single-cell spatial maps. The resulting datasets matched the transcriptomic quality and cellular composition of existing single-cell liver and kidney maps while accurately recovering the architecture of these two tissues (Extended Data Fig. 2). Notably, our method enabled high spatial mapping efficiency with over 90% of the cells mapped to a spatial coordinate (Fig. 1b).

**Figure 1:**
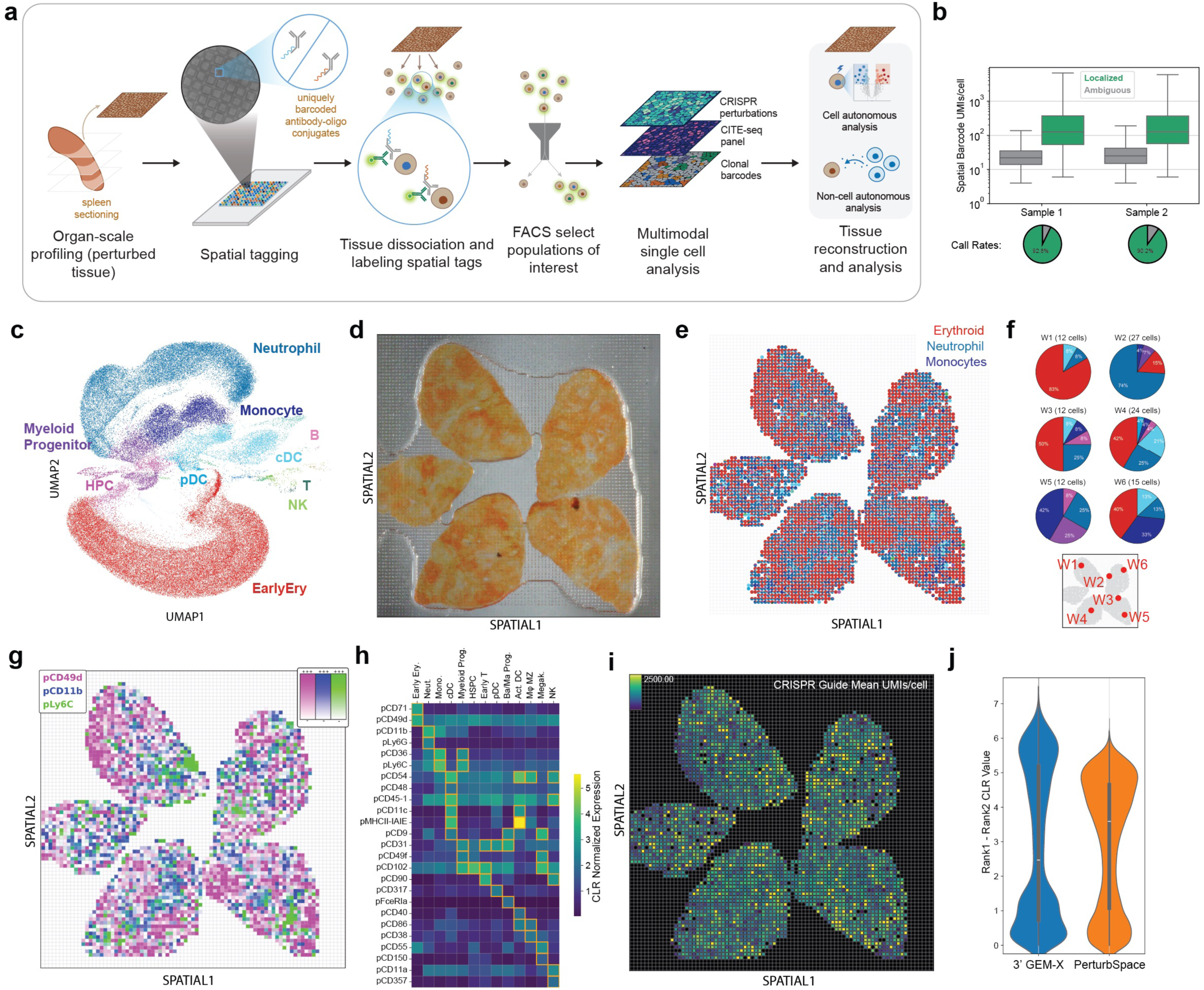
PerturbSpace enables multimodal spatial Perturb-seq at neighborhood resolution. **a**, Overview of the experimental workflow. Perturbed tissue is sectioned and tagged with spatial indexes using high-density spatial microarrays. Spatially labeled tissue sections are then dissociated into single cell suspensions and progressed through single cell workflows adapted to recover transcriptomes, spatial indexes, sgRNAs and other orthogonal genomic modalities. **b**, Spatial barcode recovery per cell. Box plots showing the number of spatial UMIs per cell for two biological samples. Cells were classified as localized (green) or ambiguous (gray). Box plots show median (center line), interquartile range (box, 25th–75th percentile), and 1.5× IQR (whiskers). Pie charts below indicate the fraction of cells with confident spatial assignments. **c**, UMAP embedding of all profiled cells in spleens colored by cell type annotation for key clusters. A full UMAP with all labels is in Extended Data Fig. 3a. **d**, Image of spleen sections on a microwell array. **e**, Spatial reconstruction of cell type composition at 80µm tissue neighborhood resolution. Cell type annotations projected onto the tissue coordinates of the spleen sections shown in c. Colors correspond to UMAP in panel c. **f**, Examples of cell type composition per splenic niche (spatial microwell). Pie charts showing the proportion of each cell type within microwells. Colors correspond to UMAP in panel c. **g**, Spatial protein expression of CD49d (erythroid), CD11b (pan-myeloid), and Ly6C (monocytic). **h,** Heatmap of CLR-normalized expression of CITE-seq antibodies across cell types. Yellow boxes highlight p<0.01, t-test. **i**, Spatial map of CRISPR sgRNA capture showing aggregated sgRNA UMIs per spatial coordinate. **j**, CRISPR sgRNA calling efficiency in PerturbSpace and Perturb-seq (3’ GEM-X) performed on the same tissue specimen.

We then evaluated the potential of our spatial barcoding approach for the multimodal analysis of in vivo perturbations. We generated perturbed splenic tissue by transplanting Cas9+ hematopoietic progenitors (HPCs) transduced with a pooled CRISPR library (88 sgRNAs) into preconditioned recipient animals. Two weeks later we profiled the perturbed spleens using our spatial hashing approach modified to simultaneously capture 4 modalities: gene expression profiles, spatial locations, surface proteomes (CITE-seq^15,16^ of 119 markers) and CRISPR perturbations (Fig. 1a, Supplemental Fig. 1). We called this approach PerturbSpace as it enables multimodal Perturb-seq at tissue neighborhood spatial resolution. PerturbSpace accurately mapped the main hematopoietic differentiation trajectories in the regenerating spleen. This includes myeloid (neutrophil, dendritic cells, and monocytes), mega-erythroid and lymphoid (B, T, and NK) lineages as well as more rare lineages such as basophils and eosinophils (Fig. 1c, Extended Data Fig. 3a). Analysis of the spatial patterns revealed the typical spleen zonation composed of a leukocytic-dense white pulp and an erythroid-dense red pulp, which indicates the regeneration of the homeostatic splenic structure (Fig. 1d-e). By zooming into specific spatial wells, we resolved neighborhoods with distinct differentiation patterns, ranging from single to multilineage niches (Fig. 1e-f). Analysis of the surface proteomes yielded high-quality, lineage-specific signals for multiple protein markers, recapitulating the red pulp versus white pulp zonation observed at the transcriptomic level (Fig. 1g-h, Extended Data Fig. 3b). Finally, we benchmarked our ability to detect CRISPR perturbations in tissue. This yielded high sgRNA detection efficiency, on par with state-of-the-art Perturb-seq methods, with 859 median sgRNA UMIs per cell and 92.5% of cells assigned to sgRNAs (Fig. 1i-j). Notably, PerturbSpace identified perturbations causing strong developmental defects in differentiating HPCs such as *Cebpa*-, *Rcor1*- and *Mbd3*-KOs (Extended Data Fig. 3c). Taken together, these results demonstrate that PerturbSpace allows for unbiased high-throughput spatial profiling of perturbed tissue across multiple modalities.

### Identifying perturbations that alter colony forming units in regenerating spleen

Building on PerturbSpace’s compatibility with high-throughput *in vivo* functional genomics, we used it to interrogate the roles of 40 transcriptional regulators during regenerative hematopoiesis in the spleen^17^. In this model, the transplanted HPCs seed the spleen and form splenic colony-forming units (CFU-S), clonal nodules whose composition reflects the developmental potential of a single founder progenitor^18^. To delineate the spatial boundaries of each individual CFU-S we built dual libraries with semirandom clonal barcodes expressed alongside 88 sgRNAs (2 sgRNAs per target gene + 8 NTCs). HPCs were isolated and expanded from Cas9 donor animals, transduced with our dual libraries and transplanted into preconditioned animals. Two weeks post-transplant we used an adapted PerturbSpace method to jointly map the transcriptomes, spatial locations, sgRNA identities and clonal barcodes of the perturbed HPC progeny across 50 sections obtained from three spleens (Fig. 2a).

**Figure 2:**
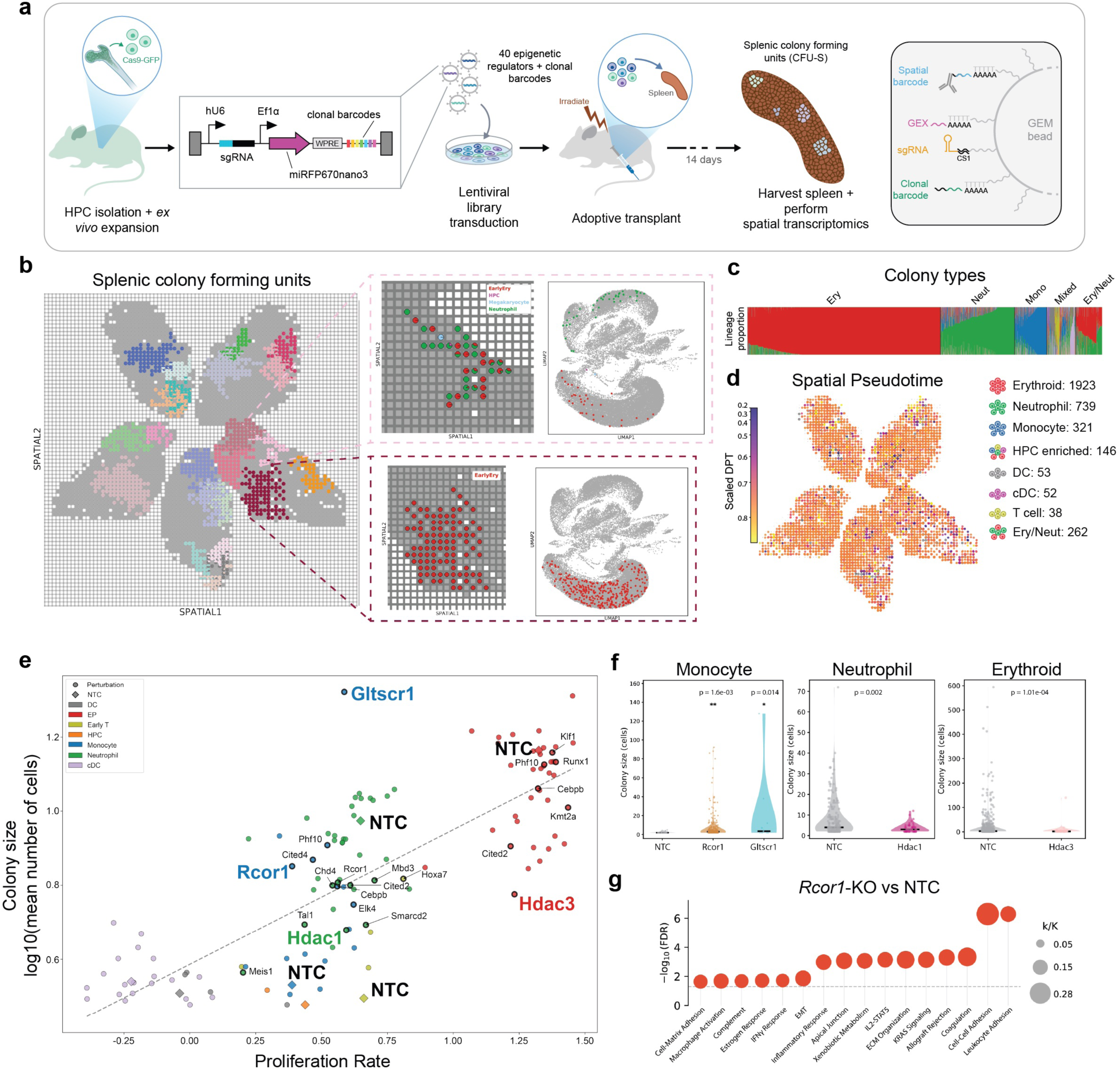
In vivo CRISPR screening of transcriptional regulators reveals perturbation-specific effects on spatial organization in the regenerating spleen. **a**, HPCs were isolated from donor Cas9 animals and expanded ex vivo, transduced with a lentiviral library targeting 40 transcriptional regulators plus clonal barcodes, and transplanted into irradiated recipient mice. After 14 days, spleens were harvested, and perturbed splenic colony-forming units (CFU-S) were profiled using PerturbSpace. b, Spatial maps of one array (80µm resolution) with select CFU-S labeled (left) with an inset showing the distribution of cells within one CFU-S. c, K-means clustering identified eight distinct CFU-S types, labeled by the predominant cell type in each cluster. Bar plot represents the proportion of all CFU-S. **d**, Spatial map of diffusion pseudotime at 80µm resolution. **e**, CFU-S size vs proliferation. Dashed line indicates linear regression fit (R² = 0.627, p = 1.11e⁻¹⁶). Perturbations with significantly different colony size compared to NTC controls of the same cell type are labelled (Mann-Whitney U test, Benjamini-Hochberg adjusted q ≤ 0.25). **f**, Size comparison between NTC and perturbed CFU-S from e. Statistical significance assessed by Mann-Whitney U test. **g**, ORA of differentially expressed genes in Extended Data Fig. 6a. Each dot represents a significantly enriched or depleted gene set (FDR < 0.05).

We obtained 342,633 high quality single cell transcriptomes with unambiguous locations at 80 µm spatial resolution. Of these, 263,863 were assigned to a specific perturbation and 179,589 contained both clonal barcodes and perturbations (Extended Data Tables 2, 3). Then, using clonal barcode and spatial tag information we were able to identify 19,174 colonies, which we defined as spatially contiguous patches of cells, with adjacent spatial tags, that share the same sgRNA and clonal barcode (Fig. 2b). Colony sizes varied from 1-163 microwells and were composed of 2-815 cells (Extended Data Fig. 4a-b, Extended Data Table 3). Overall, our colonies covered 34.4% of the tissue area of the perturbed spleens. Then, we classified the colonies larger than 5 cells based on their cell type composition using a K-means clustering approach. This identified 3,484 colonies that spanned eight distinct colony types (Fig. 2c, Extended Data Fig. 4c). The most abundant types were neutrophil- and erythroid-dominant colonies, followed by monocytic and balanced erythroid-neutrophil colonies. Multilineage, T cell-dominant, and dendritic cell-dominant colonies were the least abundant colony types (Fig. 2c, Extended Data Fig. 4c, Extended Data Table 4).

We next asked how perturbations in transcriptional regulators affect colony cell type composition, which reflects alterations in the potential of both the founder HPC and its derived progeny. Disruption of key hematopoietic transcription pioneer factors drastically changed the composition of the resulting colonies (Extended Data Fig. 4d-g, Extended Data Table 4). For instance, loss of the master myeloid regulator *Cebpa* depleted myeloid colonies (neutrophil and monocyte), yielding erythroid-only colonies. Conversely, disruption of the monocyte regulator *Irf8* ablated monocytic colonies. Other perturbations favored rare colony types: *Hoxa7*, *Hoxa9*, and *Kmt2d* enhanced T-cell colonies, while *Hdac1* promoted cDC colonies (Extended Data Fig. 4g). To complete our colony classification, we used diffusion pseudotime (DPT) to quantify the differentiation state of cells within each colony^19^ (Extended Data Fig. 5a). This enabled us to visualize colony maturation patterns *in situ* and identify perturbations that disrupt normal colony maturation (Fig. 2d, Extended Data Fig. 5b). Interestingly, we observed a spatial differentiation gradient within unperturbed colonies (NTC), where the maturation state of cells, measured by DPT, increased as their physical distance from the founder HPC grew. This finding suggests that cells are progressively displaced from the founder HPC as they mature. Supporting this model, the correlation between spatial positioning and maturation was disrupted in colonies where genetic perturbations blocked differentiation such as *Rcor1*-KO (Extended Data Fig. 5b-c).

Lastly, we examined how perturbations alter the physical size of colonies, a measurement that cannot be captured by dissociative scRNA-seq alone, where spatial relationships between clonally related cells are lost. We hypothesized that colony sizes would directly correlate with the proliferation activity of the cells within it. Our analysis confirmed this, revealing a good correlation between colony size and the proliferation score of its component cells (R^2^=0.627), a proxy for proliferation based on cell cycle gene expression (Fig. 2e, Extended Data Table 5, see Methods). Interestingly, several perturbations changed colony sizes without altering their proliferation scores, for example disruption of *Rcor1* and *Gltscr1* generated monocytic colonies significantly larger than their unperturbed counterparts (NTCs) at similar proliferation rates (Fig. 2f). We next examined the mechanisms underlying these colony size changes using differential expression analysis between perturbed and NTC colonies. *Rcor1-KO* monocytic colonies showed a marked upregulation of cell-cell and cell-matrix adhesion programs (Fig. 2g, Extended Data Tables 6, 7). This suggests that *Rcor1* ablation enhances tissue adhesion of the cells that form these colonies, which delays their shedding to the bloodstream and ultimately generates larger colonies. By contrast, disruption of *Hdac1* and *Hdac3* led to smaller neutrophilic and erythroid colonies, respectively, at similar proliferation rates (Fig. 2e-f, Extended Data Table 5). Consistent with our previous finding, *Hdac1*-KO neutrophils downregulated the expression of cell adhesion molecules (Extended Data Fig. 6, Extended Data Tables 6, 7). These findings illustrate how PerturbSpace can identify genetic programs contributing to high order tissue organization, a capability inaccessible to conventional *in vivo* Perturb-seq.

### Detection of non-cell-autonomous perturbation effects

Because PerturbSpace captures cellular neighborhoods, we reasoned it could detect non-cell-autonomous effects mediated by perturbed cells on their tissular niches. To test this concept, we chose the liver, where immune cells represent a small percentage of total cells, reasoning that using PerturbSpace we could isolate and measure the extrinsic effects mediated by perturbed immune cells on their adjacent cells. We chose overexpression of IFNγ, a strong paracrine perturbation that would produce a well characterized response in the surrounding cells. To build this model, we co-transplanted HPCs transduced with a doxycycline-inducible lentiviral vector encoding either IFNγ or a control peptide (Ctrl), each tagged with a unique expressed DNA barcode that can be detected through 3’ single-cell RNAseq. After HPC-derived immune cells colonized the liver (at 6 weeks post-transplant), we induced ORF expression for one week (Fig. 3a). Subsequently, we used PerturbSpace to characterize the spatial single cell expression profiles in perturbed liver across 4 tissue sections at 80 µm spatial resolution. This includes both ORF-expressing immune cells, derived from the transplanted HPCs, and endogenous liver cells. PerturbSpace identified 11,304 spatial single-cell transcriptomes that span all hepatic cell types including parenchymal and immune subsets (Fig. 3b-c, Extended Data Fig. 7a). The expression of IFNγ or control peptide (Ctrl) was restricted to the transplant-derived immune lineages (2496 cells) and was especially prominent in monocyte/macrophages and dendritic cells (Fig. 3d, Extended Data Fig. 7a).

**Figure 3:**
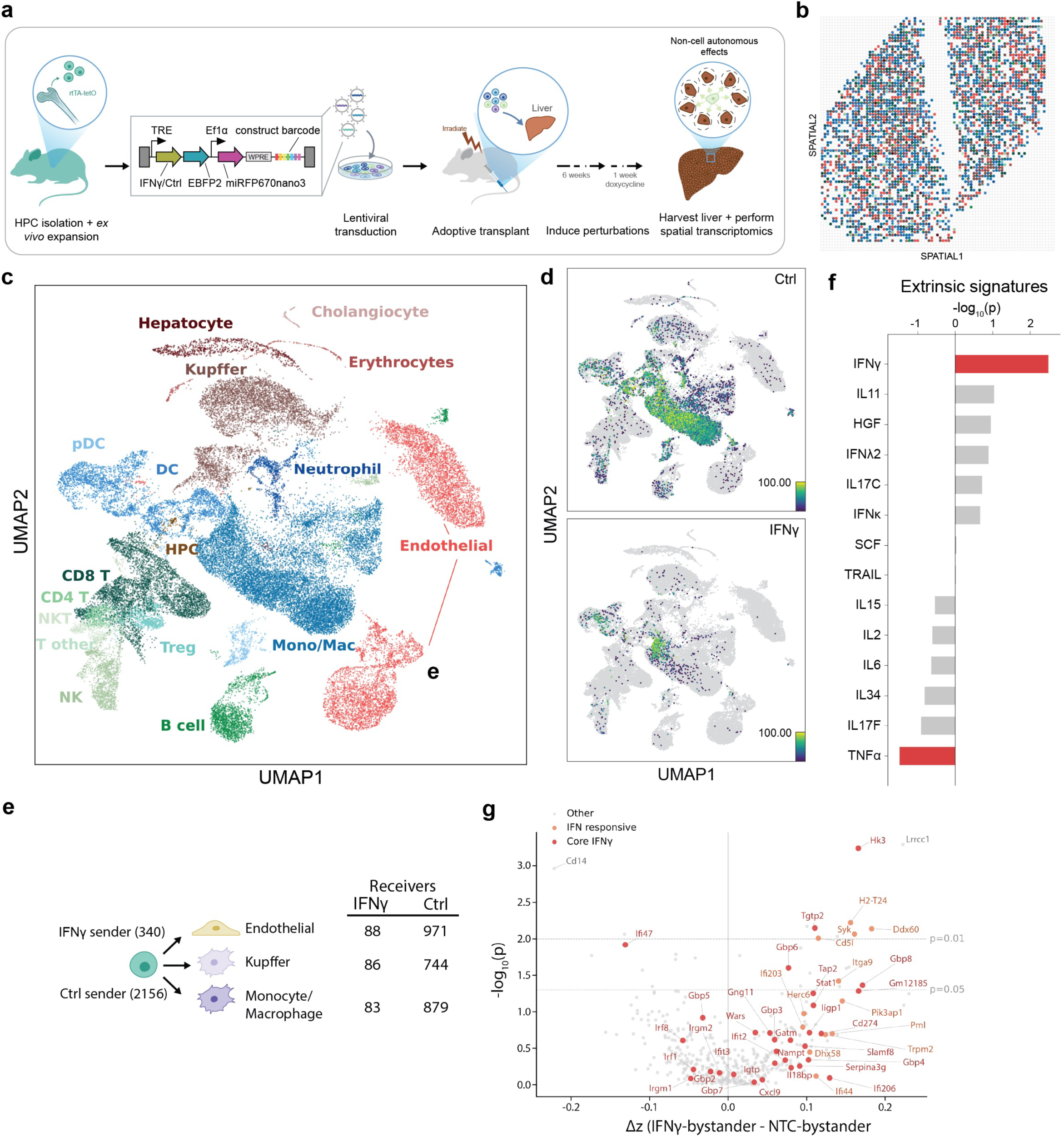
Detection of non-cell autonomous effects mediated by cytokine overexpression in the liver. **a**, Schematic of experiment. **b**, PerturbSpace array with two liver sections colored by cell type at 80µm resolution. Cell type annotations in panel c. **c**, UMAP of single cell profiles colored by cell type. Cell type colors match panel b. **d**, UMAP of single cell profiles colored by UMIs for the Ctrl and IFNγ ORF barcode. **e**, Number of IFNγ and Ctrl sender and receiver cells used for non-cell autonomous effects analysis. Neighborhood was defined as a 3 × 3 well window: IFNγ neighborhoods contained only IFNγ-overexpressing senders (UMI ≥ 10, 340 cells) and no Ctrl senders; control neighborhoods contained only Ctrl senders (UMI ≥ 10, 2,156 cells). Host cells were construct-negative cells (UMI = 0 for both constructs) of the three most abundant types in IFNγ neighborhoods: endothelial cells, Kupffer cells, and monocyte-derived macrophages (n = 257 in IFNγ, n = 2,594 in control). **f**, Change in cytokine response signatures between IFNγ and Ctrl neighborhoods. **g**, Volcano plot showing IFNγ-related gene expression changes.

To assess the effects of IFNγ secretion on neighboring cells, we performed differential expression analysis between the host cells that populated IFNγ- and Ctrl-expressing neighborhoods (Fig. 3e). We then analyzed 44 cytokine response signatures (MouSSE collection)^20^ of empirically derived response programs (see Methods, Extended Data Table 8). The IFNγ-associated signature was the most upregulated in unperturbed cells localized in IFNγ-expressing neighborhoods compared to Ctrl neighborhoods (p = 3.2 × 10⁻³) (Fig. 3f, Extended Data Fig. 7b, Extended Data Table 8). The upregulated genes in IFNγ-expressing neighborhoods included core interferon responsive genes (*Gbp6*, *Gbp8*^21^, and *Ddx60*^22^) as well as IFNγ- responsive immune pathways such as MHCI (*H2-T24*), signaling cascades (*Syk*) and transcriptional regulations (*Stat1*). Interestingly, we observed a clear blockade of IFNγ over TNF signaling in our liver model (Fig. 3g). IFNγ and TNF pathways can be either synergistic or antagonistic depending on cellular context^23^ and our data suggests an opposing behavior between these pathways in this liver model (Fig. 3f, Extended Data Table 8). These results illustrate the sensitivity of PerturbSpace to map and characterize perturbation-mediated extrinsic effects in tissues with precision.

## Discussion

In this study, we present a spatial technology that enables high throughput in vivo CRISPR screening at single-cell resolution with compatibility for multiple orthogonal proteogenomic readouts. By combining spatial cell hashing with standard single-cell sequencing workflows, we address scalability and efficiency bottlenecks, which have limited spatial functional genomics screens. Because PerturbSpace utilizes single-cell suspensions, it enables orthogonal single-cell readouts with spatial resolution which has been challenging due to assays being optimized for RNA^24^. We demonstrate this compatibility by performing spatially-resolved Perturb-seq while simultaneously measuring 119 surface proteins and tracking clonality during differentiation. This compatibility should extend in principle to epigenetic profiling (ATAC-seq^25,26^ and Cut&Tag^27^), TCR/BCR repertoire^28^, and any other readouts compatible with scRNA-seq such as metabolic labeling readouts^29^. In summary, PerturbSpace enables streamlined true multimodal spatial analysis, a capability that is not possible with nuclei-based approaches^30^.

The use of an antibody-based hashing approach confers additional advantages in experimental design. Spatially labeled cells can be enriched by FACS prior to sequencing, which is not possible with other tissue barcoding spatial technologies^30^. This capability reduces the reducing downstream library preparation and sequencing costs substantially. Additionally, FACS-based enrichment enables targeted spatial profiling of specific cell populations. This allows for the spatial mapping of selected lineages, as we demonstrate by specifically measuring perturbed- only cells in the spleen. We envision this feature will enable the spatial mapping of scarce cell types at organ scale, such as rare types of resident immune cells.

A major limitation of PerturbSpace is its relatively low resolution, which prevents the study of cellular morphology and intimate cell-cell contacts (e.g., immune synapses) in *in vivo* perturbation approaches. However, the resolution of PerturbSpace is optimal to measure spatial clonal patterns as well as non-cell-autonomous effects, which are two key processes underlying tissue dynamics and architecture. In conclusion, PerturbSpace’s ability to map both cell extrinsic and intrinsic perturbation effects at neighborhood resolution paves the way for studying the gene regulatory networks that orchestrate tissue function in homeostasis and disease^31,32^.

## Methods

### Animals

All procedures were approved by the Institutional Animal Care and Use Committee of Arc Institute (ARC-024-003). This study used C57BL/6J (Jax #000664) and Rosa26-Cas9 (Jax #028555) mice. For the non-cell autonomous experiment, we used KH2 mice (expressing rtTA from the *Gt(ROSA)26Sor* promoter) as donor mice (Jax #029415). Mixed sexes were used for donor cells and male C57BL/6J aged 8-10 weeks were used as recipient mice.

### Isolation of hematopoietic progenitors from bone marrow

Hematopoietic progenitor cells (HPCs) were isolated from the femur, tibias and pelvis from 8-12 week old donor mice (either Cas9 or KH2) by crushing the bones and filtering through a 70μm filter. Lineage negative, Ckit+ cells were enriched using MACS beads (Miltenyi #130-090-858, #130-091-224). HPCs were then expanded for 10 days in HemEx-Type9AØ (Iwai #A5P10P01C) supplemented with 10 ng/mL SCF (PeproTech) and 100 ng/mL Tpo (PeProTech).

### Cloning

For the genetic perturbation experiments, we modified the CRISP-seq backbone (Plasmid #85707) by replacing the eBFP2 reporter by miRFP670nano3^33^ and by inserting a CS1 capture sequence in the CR1 scaffold^34^. Clonal barcodes were designed as WSNNNWSNNNWSNNNWSNNNWSNNNWSNNNWSNNNW^35^, ordered as a single stranded DNA library (IDT). CRISPR and clonal barcodes were cloned in sequential steps using NEB HiFi DNA Assembly Master Mix. A list of gRNA sequences can be found in Extended Data Table 1. For the cytokine overexpression experiment, we designed a lentiviral backbone comprising an inducible TRE promoter driving cytokine ORF expression followed by a P2a peptide and eBFP2 reporter, a constitutive EF1a promoter driving miRFP670nano3. The backbones were ordered from VectorBuilder.

### Lentiviral production

We produced VSV G pseudotyped lentivirus using pMD2.G (Plasmid #12259) and psPAX2 (Plasmid #12260), and Lipofectamine 3000 (ThermoFisher) with Lenti-X 293T cells (Takara, 632180). Two days after transfection the supernatant was filtered at 0.45μm and concentrated 50X using a 100kDa Amicon centrifugal filter (Millipore).

### Spatial array fabrication and filling

The microwell array-based spatial hashing approach was developed by Survey Genomics (surveygenomics.com). The microwell array chips were fabricated using different processes depending on their feature sizes. Chips were designed to be microscope slide-sized and versioned according to various well dimensions. The microwell arrays were fabricated using PDMS soft lithography from a custom patterned wafer (Celldom) and bonded to microscope slides using super glue. Once fabricated, the chips were then filled via piezo-driven non-contact liquid dispensing (sciFLEXARRAYER S3, Scienion AG) using a custom program to fill the wells with a specific barcode layout.

### Spatial labelling mixture design (Antibody-oligo conjugate)

To enable spatial barcode labeling of whole cells in situ, we used a cocktail containing *a*MHC-I and *a*CD45 antibodies designed to stain nucleated mouse cell types (TotalSeq-B Hashtag 1-24, BioLegend). Each antibody is conjugated with an oligo containing a PCR handle, a barcode sequence, and a CS1 binding sequence. The CS1 binding sequence is used to hybridize a second oligo to the antibody oligo via a reverse complementary sequence CS1. The hybridized oligo contains a spatial barcode denoting a coordinate location and a poly(A). The poly(A) sequence hybridizes to the poly dT capture of the 10x Genomics 3’ GEM-X bead. Hybridization of a unique oligo to a common sequence allows us to avoid conjugating hundreds of AOCs, therefore significantly reducing costs. The oligo directly conjugated to the antibodies of the cocktail is still recoverable if desired (i.e. for chip multiplexing).

### Spatial hashing of tissue sections

Tissue sectioning was performed with a Precisionary Compresstome. Sections were first mounted on a glass slide, positioned on top of the spatial array, and clamped. Sections undergoing spatial hashing were incubated in a humidified chamber at room temperature for 30 minutes. Following the incubations sections are released from the array, briefly washed in PBS, and transferred to collagenase at 10C for dissociation. Dissociated cells were then incubated in an anti-rat PE secondary antibody (BioLegend, 405406) to label spatial hashing antibodies and sorted for 10x 3’ scRNA-seq profiling.

### Multimodal PerturbSpace

Spatially hashed cells were encapsulated in droplets using the 10x Genomics 3’ GEM-X v4 kit with adapted protocols to profile the following modalities (Supplemental Fig. 1):

- <u>Transcriptome (scRNA-seq).</u> After cDNA amplification cleanup, Single-cell transcriptomes were generated from the bead-bound cDNA following the standard 3’ mRNA 10X Genomics workflow.
- <u>Spatial modality.</u> Spatial tags were recovered as described by spiking the Spatial additive primer (5’-CCTTGGCACCCGAGAATT*C*C-3’) into the cDNA amplification mix. After cDNA cleanup, the spatial tags were dialed-out from the supernatant and indexed using custom i7 (e.g., 5’-CAAGCAGAAGACGGCATACGAGATAGGCGACTAGGTGACTGGAGTTCCTTGGCAC CCGAGAATTC*C*A-3’) and i5 (e.g., 5’-AATGATACGGCGACCACCGAGATCTACACCGAACGATGGACACTCTTTCCCTACAC GACGC*T*C-3’) primers.
- <u>Surface proteome modality.</u> After tissue dissociation the splenic cell suspension were incubated with the TotalSeq-A Mouse Universal Cocktail (BioLegend, 199901) following manufacturer’s instructions. Then, after cDNA cleanup the TotalSeq-A antibody indexes were dialed-out from the supernatant using the same primers as the spatial tags.
- <u>CRISPR modality.</u> After cDNA cleanup, the sgRNA sequences were dialed-out from the supernatant using Feature cDNA Primers 2 following the manufacturer’s protocol (10X Genomics). The resulting PCR product was amplified using Feature SI Primers 3 and indexed using the Dual Index Plate NT.
- <u>Clonal Barcode modality.</u> Following GEM-RT cleanup, a Clonal barcode primer (CBC-preamp: 5’-TCCAGCGGACCTTCCTT-3’) was spiked into the cDNA amplification mix. This primer binds to the WPRE sequenced, located upstream of the semi-random sequence that define clonal identifiers. This enables to increase the capture efficiency of the cDNA molecules containing clonal information during cDNA amplification. After cDNA cleanup. The clonal barcode sequences are dialed-out from the supernatant using a Read2-WPRE (5’-GTGACTGGAGTTCAGACGTGTGCTCTTCCGATCTCTCGACGGATCTCATGCTCGAG-3’) and a Read1 primer (5’-ACACTCTTTCCCTACACGACGCTC-3’) and indexed with 10x Genomics Dual Index Plate TT.

### PerturbSpace in the spleen (Colony forming unit analysis)

Expanded Rosa26-Cas9 HPCs were transduced with lentiviral particles at a MOI of 0.2. Transduced cells were cultured for 5 days and sgRNA-containing HPCs were sorted for the presence of miRFP670nano3 on a Highway1 Cell Sorter (STEMCELL Technologies). Three recipient mice (C57BL/6J) were irradiated twice with 4.5 Gy irradiation, 8 hours apart using a Kimtron Biological Irradiator SC500 and 300,000 sorted HPCs (miRFP670nano3+) were injected into the tail vein. Two weeks later, mice were euthanized and the spleens dissected and spatially hashed as described above. Then, spatially-hashed cells carrying genetic perturbations (PE+, miRFP670nano3+) were processed for PerturbSpace using the multimodal workflow described above to capture transcriptomes, spatial tags, surface proteomes, sgRNAs and clonal barcodes. In total 50 spleen sections and 10 arrays were used for the organ-scale study (Extended Data Table 2).

### PerturbSpace in the liver (Cytokine overexpression analysis)

Expanded KH2 HPCs were transduced with lentiviral particles at a MOI of 0.2. Transduced cells were cultured for 5 days and library-containing HPCs were sorted for the presence of miRFP670nano3 on a Highway1 Cell Sorter (STEMCELL Technologies). C57BL/6J mice were irradiated twice with 4.5 Gy irradiation, 8 hours apart using a Kimtron Biological Irradiator SC500 and 500,000 transduced HPCs were injected into the tail vein. Six weeks later, doxycycline (50 μg/g) was administered to the transplanted animals intraperitoneally daily for one week to induce cytokine expression. Lastly, animals were euthanized, livers were perfused with PBS to wash circulating immune cells and spatially hashed as described above. Then, all spatially hashed cells (PE+) were processed for PerturbSpace to capture transcriptomes, spatial tags, and ORF barcodes. In total, we spatially labelled 4 sections, with 2 sections per array

### Sequencing

All libraries were sequenced with 28 cycles for R1, 90 cycles for R2 and 10 cycles for each index. We aimed for 600M reads for gene expression libraries, 50M reads for CRISPR libraries, 100M reads for spatial barcode libraries, and 100M reads for clonal barcode libraries. All sequencing was performed on an Illumina NovaSeq 6000.

### Data processing and primary analysis

Data processing and primary analysis was performed on an Elastic Compute Cloud instance type m5n.8xlarge (Amazon Web Services). BCLs were demultiplexed to FASTQ files using bcl2fastq2 with default parameters. FASTQs were counted and aligned using cellranger (v8.0.1 and v10.0.0) using the multi subcommand ^36^. Counting and alignment of all non-transcriptomic libraries were run in Antibody Capture mode with custom feature reference files and always run in conjunction with the transcriptome in Gene Expression mode to enable accurate cell barcode counting. Because the TotalSeq-A Universal Cocktail and spatial barcode libraries are amplified with identical primer pairs, this library was aligned to a combined reference of spatial barcode and protein expression antibody barcode sequences. Because the clonal barcode libraries contained degenerate barcodes, the feature reference for each sample was created from the FASTQs themselves using custom scripts that scan the Read 2 files for the barcodes of the form WS–(NNNWS)⨉7, combine across single-cell reactions, and collapse into a consensus list with edit distance 2.

The resulting gene x feature matrices from the Cell Ranger outputs were further processed in Python using Jupyter notebooks (Python 3.12.12) and the scanpy (1.10.4) single-cell analysis package ^37^. Matrices for each library/modality and each sample were combined into a single data object with mudata (0.3.2) ^38^, merging on the filtered cell barcodes output from the spatial barcode counting and alignment. Genes were filtered to those expressed with at least 10 counts and detected (counts > 0) in at least 20 cells. PCA was run on the resulting matrix, followed by Harmony batch correction (harmonypy, 0.2.0) ^39^ and UMAP dimensionality reduction. Leiden community detection was used to identify cell clusters with similar gene expression which were then manually annotated. Cell type annotations were done based on canonical lineage markers, assigning each Leiden cluster manually to a cell type based on signature expression (see Extended Data Table 1 for markers and cluster assignments). Per-sample sequencing and QC metrics are listed in Extended Data Table 2.

### Spatial barcode calling

A combinatorial spatial barcode of X and Y (columns and rows) was used to assign individual cells to microwell positions (Extended Data Table 1). To assign cells to locations, we applied a stepwise algorithm that infers each cell’s position from the specificity of its spatial barcode signal. Cells with fewer than 5 counts were excluded as low-confidence. For each spatial index, cells with a ratio ≥ 2 between the 1st- and 2nd-ranked barcode were assigned to the 1st-ranked barcode. Remaining cells, where this ratio threshold was not met for one or both indices, were resolved using a distance- and rank-weighted scoring of combinatorial barcodes, with each cell assigned to the highest-scoring valid X–Y combination.

### Algorithm: Spatial barcode assignment

Input: cell × barcode count matrix, spatial layout, *max_dist* = 250 µm, *r* = 0.8, *d* = 0.2

1. Initialize all cells as UNASSIGNED
2. For each cell with total barcode counts below threshold:

— Assign AMBIGUOUS
3. For each cell where the top barcode exceeds ratio threshold:

— Assign top-ranked barcode
4. For each remaining UNASSIGNED cell:

— Retrieve top-n candidate barcodes ranked by count
— Generate all pairwise combinations of candidates
— Discard pairs not present in the spatial layout
— For each valid pair within *max_dist*:

— Compute score = *r* × rank_score + *d* × distance_score
— If valid pairs exist:

— Select the highest-scoring pair
— Assign the barcode with the better rank from that pair
— Else:

— Assign AMBIGUOUS

Output: per-cell spatial barcode assignments

Dispensing errors can cause certain spatial barcodes to be over-represented, leading to spurious labeling of cells from neighboring wells. To correct for these filling artifacts and assuming spatial barcode frequency is locally compositional, we normalize barcodes whose counts exceed the median counts of that spatial index (i.e. X or Y, for row-column-formatted chips) by *p* = 30% of the median. To keep the total counts per cell constant (i.e. compositional), we assign barcode UMIs from over-represented barcodes (*C*_*i*_) to neighboring under-represented barcodes (whose count ratio 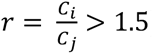) according to an adjacency tensor *A*, where *A*_*d*,*i*,j_represents the proportion of barcode 1 among the neighbors of barcode *i* at a distance *d*, iterating until there are no over-represented barcodes. When assessed on all available data, we determined that the parameters (*p* = 0.3, *r* = 15) reduced the frequency of cells assigned to over-represented barcodes by localizing outside tissue areas to background levels, consistent with all other barcodes.

Tissue sections placed on the array were imaged using a wide-field benchtop digital microscope. These images were manually edited using a photo editor to correct for fish-eye distortions and align the array to its digital representation. Once images were aligned, cellular data was able to be superimposed on the images, and a custom segmentation GUI was used to annotate wells on the chip that were overlaid with tissue

### CRISPR guide and clonal barcode calling

CRISPR guide identities and clonal barcode identities were assigned to individual cells using Geomux^40^. Guide calling and barcode calling were performed independently by passing the respective cell × feature UMI count matrices through the Geomux pipeline, briefly described below.

For each modality, cells with fewer than 5 total UMIs and features with fewer than 10 total UMIs were excluded. To correct for ambient contamination, one count was subtracted from every nonzero entry and resulting zeros were removed. Feature enrichment within each cell was then assessed using a one-sided hypergeometric test, where the observed UMI count for a given cell–feature pair was tested against the expected distribution given the total UMIs in the matrix, the total UMIs for that feature, and the total UMIs for that cell. P-values were corrected for multiple testing using the Benjamini–Hochberg procedure, and pairs with FDR ≤ 0.05 were flagged as significant. To remove borderline assignments, a log odds ratio (LOR) filter was applied using an adaptive threshold scaled to the mean nonzero UMI count in the corrected matrix. Within each cell, significant assignments were evaluated in order of decreasing significance, and those with LOR below the threshold were revoked. Each cell received zero, one, or multiple feature assignments along with corresponding UMI counts, FDR values, and LOR values.

### Diffusion Pseudotime (DPT) per cell

Diffusion pseudotime was computed on non-targeting control (NTC) cells to establish a reference differentiation trajectory. NTC cells were filtered to progenitor, erythroid, neutrophil, and monocyte populations, excluding doublets and high-mitochondrial cells. Gene expression values were log-normalized, subsetted to highly variable genes (minimum mean 0.0125, maximum mean 3, minimum dispersion 0.5), and scaled with a maximum value of 10. Principal component analysis was performed (50 components, ARPACK solver), followed by construction of a k-nearest-neighbor graph (k = 15, 50 PCs) and computation of a diffusion map with 15 components. DPT was then computed using 3 diffusion components. The root cell was selected as the HSC closest to the median position of all HSCs in diffusion component space (components 1–3), providing a stable starting point for the pseudotime axis.

To assign pseudotime to perturbed cells, PCA was recomputed on combined NTC and perturbed cells using the same preprocessing and cell type filters, excluding cells with multiple or no guide assignments. For each perturbed cell, the 15 nearest NTC neighbors were identified and pseudotime was assigned as the mean DPT of those neighbors.

To enable cross-lineage comparison, DPT values were scaled per cell type by dividing by the maximum DPT observed within each lineage branch (erythroid progenitors, neutrophils, monocytes), rescaling pseudotime to the range [0, 1]. Common progenitors retained unscaled values. Scaled pseudotime was used in all subsequent spatial analyses of differentiation state.

### Proliferation score

Cell cycle activity was scored using a cell cycle gene set^41^. S-phase and G2M-phase scores were computed using scanpy’s ‘score_genes_cell_cyclè function on the normalized log₁p-transformed expression. For each cell, S-phase and G2M-phase scores were calculated as the mean expression of phase-specific gene sets minus the mean expression of control genes matched for expression level. A per-cell proliferation score was defined as the sum of the S-phase and G2M-phase scores, providing a single continuous measure of cell cycle activity.

### Colony Identification

Upon plotting clonal barcode expression spatially, we noted that clonal barcodes typically showed high expression localized to specific areas of the chip, suggesting clonal expansion. To extract the specific microwells of each pocket of expression in a standardized way, we used a simple object detection algorithm based on image processing. Briefly, for each chip, we plotted a spatial map of the total counts of each clonal barcode per microwell with at least 3 cells. Expression values for wells without sufficient cell numbers were inferred using linear interpolation, and the resulting dense expression maps were smoothed using a Gaussian filter (σ = 1.0) to merge nearby fragmented signals. Spatial colonies were then identified by thresholding the resulting smoothed map (threshold = 1.5) and grouping adjacent thresholded regions via connected component labeling. From these candidate colonies, we further filtered out those whose total area < 9 wells or if the proportion of interpolated wells exceeded 50%.

Cells localized within the identified spatial boundaries were then evaluated for clonal barcode purity. Colony assignments required a minimum count threshold (≥10 UMIs) and a 1st-to-2nd ratio ≥ 2.0 (similar to what was done for spatial barcodes), with cells failing these criteria flagged as ambiguous. Consensus clonal identities were assigned to each spatial colony if a specific lineage represented at least 30% of the confidently assigned cells within the pocket. These colonies formed the initial seed CFU-S, that were then refined as described below.

### Colony Refinement

CFU-S were defined as spatially contiguous groups of cells sharing both a clonal barcode identity and a CRISPR guide identity. Starting from seed colony definitions (described above), we applied a multi-step refinement pipeline, taking barcode into account, independently to each chip.

Spatially profiled cells were assigned to seed colonies if their array position was occupied by that colony and their guide assignment exactly matched the colony’s guide identity. Unassigned cells were then subjected to de novo colony discovery: cells sharing identical section, barcode, and guide identities were grouped, and spatial adjacency graphs were constructed using an 8-connected Moore neighborhood (cells in wells differing by at most one position in row and column). Connected components of two or more cells were designated as new colonies; singletons were discarded.

Colonies were then refined through sequential steps. First, within-colony barcode heterogeneity was resolved by designating the most frequent barcode as the dominant identity and splitting cells carrying alternative barcodes into sub-colonies. Colonies that became spatially non-contiguous after barcode splitting were further partitioned into connected components. Next, unassigned cells were iteratively annexed into adjacent colonies if their guide and barcode identities exactly matched the colony’s modal identities, with all eligible annexations applied simultaneously per iteration. Adjacent colonies within the same tissue section sharing identical guide and barcode identities were then merged iteratively.

Post-refinement cleanup included removal of cells whose barcodes did not overlap the colony’s founding barcode set, re-application of contiguity splitting and merging, dissolution of colonies lacking any barcoded cell, removal of singleton colonies, and a final pass of de novo discovery to recover remaining unassigned clusters. Per-colony attributes for all 19,174 CFU-S are provided in Extended Data Table 3.

Colony assignments were validated through a comprehensive test suite assessing identity coherence (unique guide and barcode sets form connected components, single-guide cells target exactly one gene), spatial integrity (colonies form single connected components with articulation point fraction not exceeding 30%), assignment exhaustion (no unassigned cell with matching barcode and guide neighbors remains), and internal consistency (single section per colony, minimum size of two, no multi-colony assignments).

### Colony clustering by composition

For each colony, a cell type frequency vector was computed. Doublets, high-mitochondrial cells, fibroblasts, and endothelial cells were excluded from frequency vectors. Colonies with fewer than 5 cells were removed.

The optimal number of clusters was determined using the elbow method applied to the within-cluster sum of squares across K = 4–15, selecting K at the point of maximum second derivative of the inertia curve, which yielded K = 5. K-means clustering (20 random initializations) was performed on the colony cell type frequency matrix.

Perturbation enrichment within each cluster was assessed using two-sided Fisher’s exact tests on 2 × 2 contingency tables comparing colony counts for each perturbation inside versus outside the cluster. Perturbations with fewer than 2 colonies were excluded. P-values were corrected using the Benjamini–Hochberg procedure (FDR ≤ 0.05), and associations were filtered to require |log₂(odds ratio)| ≥ 0.5.

To resolve heterogeneity within one cluster (C3), a second round of K-means clustering was performed on the subset of C3 colonies using the same elbow method (K = 4) and enrichment testing procedure. K-means centroids and per-perturbation cluster enrichment are reported in Extended Data Table 4.

### Colony size vs. proliferation

Each CFU-S was annotated with its majority cell type, defined as the cell type comprising the highest fraction of cells. To restrict analysis to colonies with clear single-lineage identity, only colonies in which the majority cell type accounted for at least 75% of cells were retained. Colonies were further required to contain at least 2 cells, have a valid perturbation assignment, and have a non-missing proliferation score. Doublets, high-mitochondrial cells, fibroblasts, and endothelial cells were excluded, and cell type-perturbation combinations represented by fewer than 5 colonies were removed.

Colony proliferation score was defined as the maximum per-cell proliferation score within the colony. Colony size was measured as the number of cells per colony. For each perturbation-cell type combination, colony sizes were compared to the corresponding NTC colonies using two-sided Mann–Whitney U tests, with p-values corrected using the Benjamini–Hochberg procedure (FDR ≤ 0.25).

To assess the relationship between proliferation and colony size, colonies were grouped by cell type-perturbation combination and the mean proliferation score and mean colony size were computed. A linear regression was fit on these aggregated values (mean proliferation score versus log₁₀ mean colony size) and the coefficient of determination and F-test p-value were reported. Per (perturbation × cell type) Mann–Whitney statistics and the overall regression fit are reported in Extended Data Table 5.

### CFU-S effective cell type number

Colony cell type diversity was quantified as the effective number of cell types, defined as exp(*H*) where *H* = −Σ pᵢ ln pᵢ is the Shannon entropy and pᵢ is the proportion of cell type i. This metric represents the number of equally abundant cell types that would produce the observed entropy.

Colonies were filtered to those with at least 5 cells and a valid perturbation assignment. Because colony diversity is expected to increase with size, we tested whether perturbations alter diversity beyond what colony size alone would predict. For each perturbation with at least 50 colonies, perturbation and NTC colonies were pooled and an ordinary least-squares model was fit with effective number of cell types as the response variable and perturbation status and log₁₀ colony size as predictors. The significance of the perturbation coefficient was used to assess whether diversity differed after controlling for colony size. P-values were corrected across perturbations using the Benjamini–Hochberg procedure (FDR ≤ 0.05). Per-perturbation diversity results are listed in Extended Data Table 4.

### CFU-S DPT as function of distance from CFU-S HSC

To examine the spatial relationship between differentiation state and proximity to founder HSCs, analysis was restricted to colonies containing at least one HSC and at least 5 cells. For each colony, the Euclidean distance from every non-HSC cell to the nearest HSC within the same colony was computed from spatial well coordinates and paired with the cell’s scaled DPT value.

The relationship between differentiation state and HSC distance was characterized using locally weighted scatterplot smoothing (LOWESS, smoothing fraction 0.4) applied separately per perturbation. 95% confidence intervals were estimated from 200 bootstrap resamples. As a complementary parametric summary, ordinary least-squares regression of scaled DPT on distance to nearest HSC was fit per perturbation, and the slope, R², and F-test p-value were reported.

### Scaled DPT comparison per cell type and perturbation

Per-colony scaled diffusion pseudotime (DPT) was computed as the mean scaled DPT across cells of the colony’s majority cell type (ct2). Colonies were retained if they had a majority cell-type purity ≥ 75 %, at least one majority-cell-type cell with a defined scaled-DPT value. For each cell type we then identified candidate perturbations as those with ≥ 4 colonies. For each perturbation, the colony-level scaled-DPT distribution was compared to the matched NTC distribution from the same cell type using a two-sided Mann–Whitney U test. Within each cell-type panel, raw p-values across all tested candidates were corrected for multiple testing with the Benjamini–Hochberg (BH) procedure.

### DGE and ORA between NTC and perturbed CFU-S in spleen

Differential gene expression between perturbed and NTC CFU-S was assessed using a pseudobulk approach. For each cell type-perturbation combination in which colony size significantly differed from NTC, colonies dominated by that cell type (≥75% of cells) with at least 2 cells were selected. Raw UMI counts were summed across all cells of the target cell type within each colony to produce one pseudobulk profile per colony, requiring at least 2 colonies per group.

Differential expression was tested using limma-trend (3.66.0, (R 4.5.2)). Moderated t-statistics and Benjamini–Hochberg adjusted p-values were obtained from the perturbation–NTC contrast. For cases where too few colonies were available for colony-level replication, pseudobulk profiles were instead constructed per chip, with each chip serving as an independent replicate using the same limma-trend approach.

Over-representation analysis (ORA) was performed on significantly upregulated and downregulated gene lists separately using a curated collection of 73 gene sets comprising MSigDB Hallmark pathways (v7.5), MSigDB Canonical Pathways, and Gene Ontology Biological Process terms manually selected for relevance to hematopoiesis, myeloid differentiation, immune signaling, chromatin regulation, cell adhesion, and cell death. Human gene symbols were converted to mouse orthologs, and gene sets with fewer than 10 or more than 500 members were excluded. Enrichment was assessed using one-sided Fisher’s exact tests against the background of all genes tested in each comparison, with Benjamini–Hochberg correction applied across all testable gene sets. Pseudobulk-level limma-trend results for each contrast are provided in Extended Data Table 6; the curated gene sets and per-contrast ORA results are provided in Extended Data Table 7.

### Non-cell-autonomous (NCA) IFNγ paracrine analysis

For each cell, the cell was defined as a construct cell based on a geomux call and retained as a sender of construct X (X ∈ {IFNγ, NTC}) only if its construct-X UMI count exceeded 10. Construct-positive cells failing this UMI threshold were marked LowExpr and excluded from the sender pool.

Each cell was assigned to a well via its (chip, array-row, array-column) coordinates. For every well we counted effective senders over its 3×3 grid neighborhood: the well plus its eight immediate neighbors on the same chip. A well was classified IFNγ-pure if its neighborhood contained at least one IFNγ sender and zero NTC senders, and symmetrically NTC-pure if it contained at least one NTC sender and zero IFNγ senders. All other wells were left unassigned.

Focal cells were defined as cells with no construct (based on geomux) after removing low-quality classes (such as doublets). Each focal cell was tagged according to the purity classification of its well as an IFNγ-bystander, NTC-bystander, or unassigned. The analysis was restricted to focal cells from the top three most abundant cell types: Kupffer, Mono/Mac, and Endothelial, which were jointly designated the focal receiver group. Per-cell gene expression counts in the focal pool were normalized to 10⁴ total UMIs and log1p-transformed.

Per-cell cytokine activity scores were estimated using the mouse cytokine response signatures from the MouSSE gene-set network^20^, which derives its signatures from the Immune Dictionary^42^. Scores were computed using the multivariate linear model (mlm) implemented in decoupler (2.1.6). Prior to scoring, each MouSSE signature was restricted to target genes detected in ≥15% of focal cells of the focal receiver group; the same expression filter was applied identically across all signatures to preserve cross-cytokine comparability. Per-cell activity scores were then z-scored within each focal cell type (computed separately for Kupffer, Mono/Mac, and Endothelial), with the reference distribution for each (cytokine, cell type) consisting of all focal cells of that cell type that were not IFNγ-bystanders.

The IFNγ paracrine effect was assessed by a two-sided Mann–Whitney U test on within-cell-type z-scored MouSSE-IFNγ activity, comparing IFNγ-bystander to NTC-bystander focal cells of the focal receiver group. The Mann–Whitney test was then repeated independently for each of the MouSSE cytokine signatures. Cytokines were ranked by signed −log₁₀(p) (sign of the mean Δz between IFNγ- and NTC-bystander cells), with Benjamini–Hochberg FDR correction applied across the cytokines.

The Gene-level paracrine effects for each gene in the MouSSE-universe were assessed (n = 417): per-cell expression was z-scored within focal cell type, and the difference of mean z-scores between IFNγ-bystander (n = 257) and NTC-bystander (n = 2,594) focal cells was computed (Δz). Per-gene significance was assessed by a two-sided Mann–Whitney U test on the per-cell z-scores.

The cytokine response signatures used, and the per-cytokine results are listed in Extended Data Table 8 (sheets MouSSE_signatures and cytokine_NCA_results); per-cell focal IFNγ z-scores and per-cell scores for all 44 cytokine signatures are also provided in Extended Data Table 8. Per-gene results for the IFNγ-bystander vs Ctrl-bystander comparison are reported in Extended Data Table 8 (sheet IFNg_neighbour_DGE).

## Acknowledgements

We thank Gabriel Grenot for help with animal work. We thank Josh Mast, Alyssa Ward, and Xiaoying Meng for assistance with experiments. We thank Alexander Marson, Sean Scott and Brian Plosky for comments on the manuscript. AAN, CDV, KR, JJB, MEC, AD, CR-T, ML, IA, and DL-A are supported by Arc Institute.

## Author contributions

AAN and DL-A conceived the project and designed the PerturbSpace workflow. DL-A, CDV and AAN conceived and developed the spatial clonal assay. AAN, CDV, KR, JJB developed the in vivo perturbation assays and performed the experiments. GCH and HL developed the microwell array and dispensing procedures. AAN, KR, JJB, and HL performed the PerturbSpace profiling. JB and LV developed the computational framework for clonal barcode analysis. GCH processed and annotated PerturbSpace datasets and performed the analysis of the orthogonal modalities with input from IA, MECC, AAN, CJY, and DL-A. IA analyzed the PerturbSpace data and extracted biological insights with input from AD, ML, CJY, AAN, and DL-A. AAN, CR-T, and DL-A designed figures. IA and DL-A supervised the project. AAN and DL-A wrote the manuscript with input from IA.

## Data and Code availability

A Python package to analyze Survey Genomics spatial data is available at https://github.com/survey-genomics/survey/. Raw data and custom code will be available upon publication.

---

**Extended Data Figure 1:**
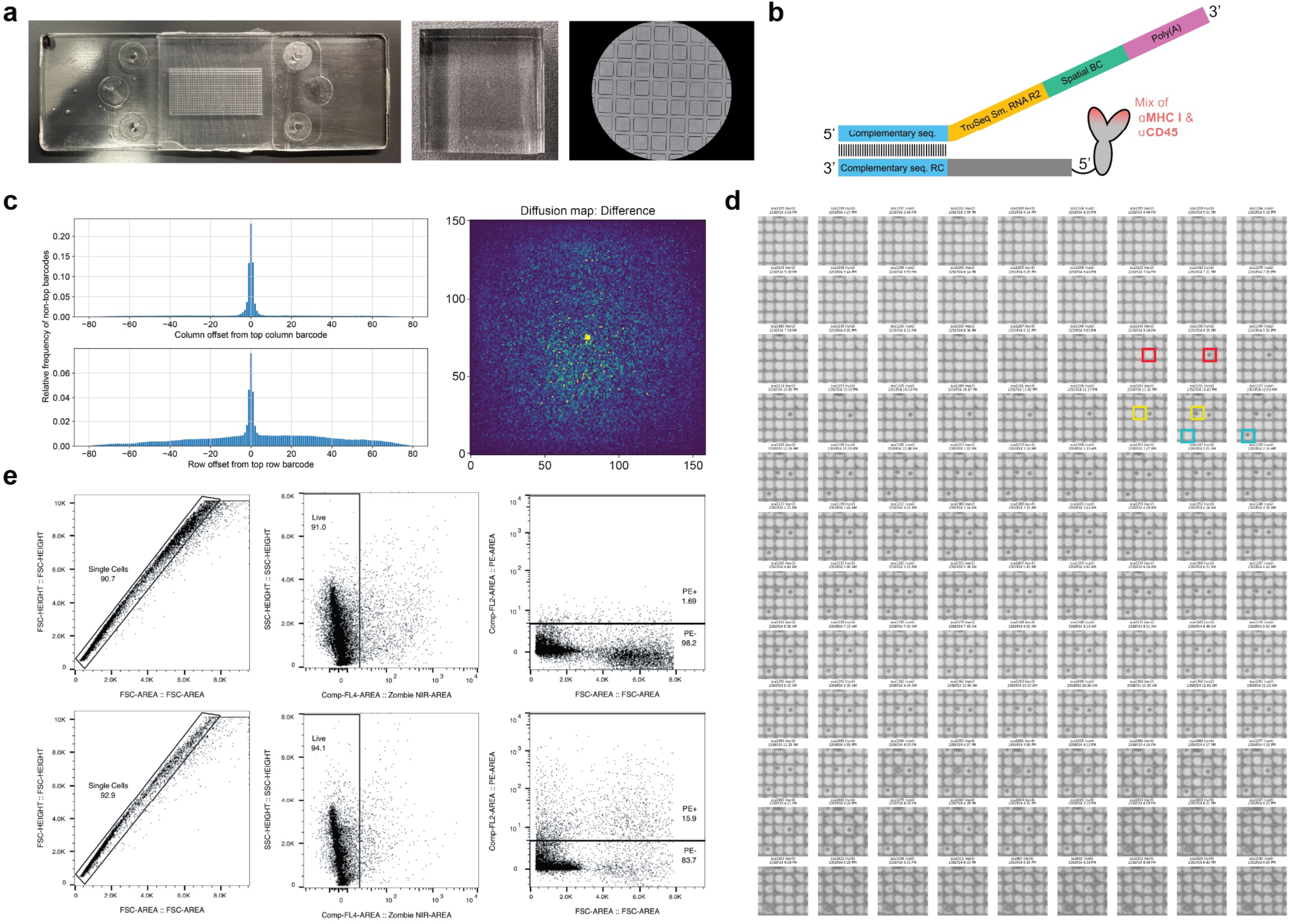
Design of PerturbSpace assay. **a**, Images of the spatial microwell chip. **b**, Schematic of the spatial oligo hashtag. Spatial hashing oligos are hybridized to anti-CD45 and MHCI TotalSeq-B Hashtag antibodies by dispensing the antibody mix into the pre-indexed spatial microarray. Spatial oligos are then captured during the RT step in 3’ scRNA-seq. **c**, Diffusion profile of spatial barcodes. Left: Histograms showing the relative frequency of non-top barcodes as a function of column offset (top) and row offset (bottom) from the top-ranked barcode. Right: diffusion map showing the spatial distribution of the spatial barcode. **d**, Time-lapse images of deposition of antibodies and spatial oligos in microwells. Colored boxes highlight three deposition examples. **e**, FACS enrichment of spatially tagged cells using anti-spatial tag PE secondary antibody. Top row is background control for PE. Bottom row corresponds to one of the spatially hashed spleen sections that generated the results in Fig. 1b.

**Extended Data Figure 2:**
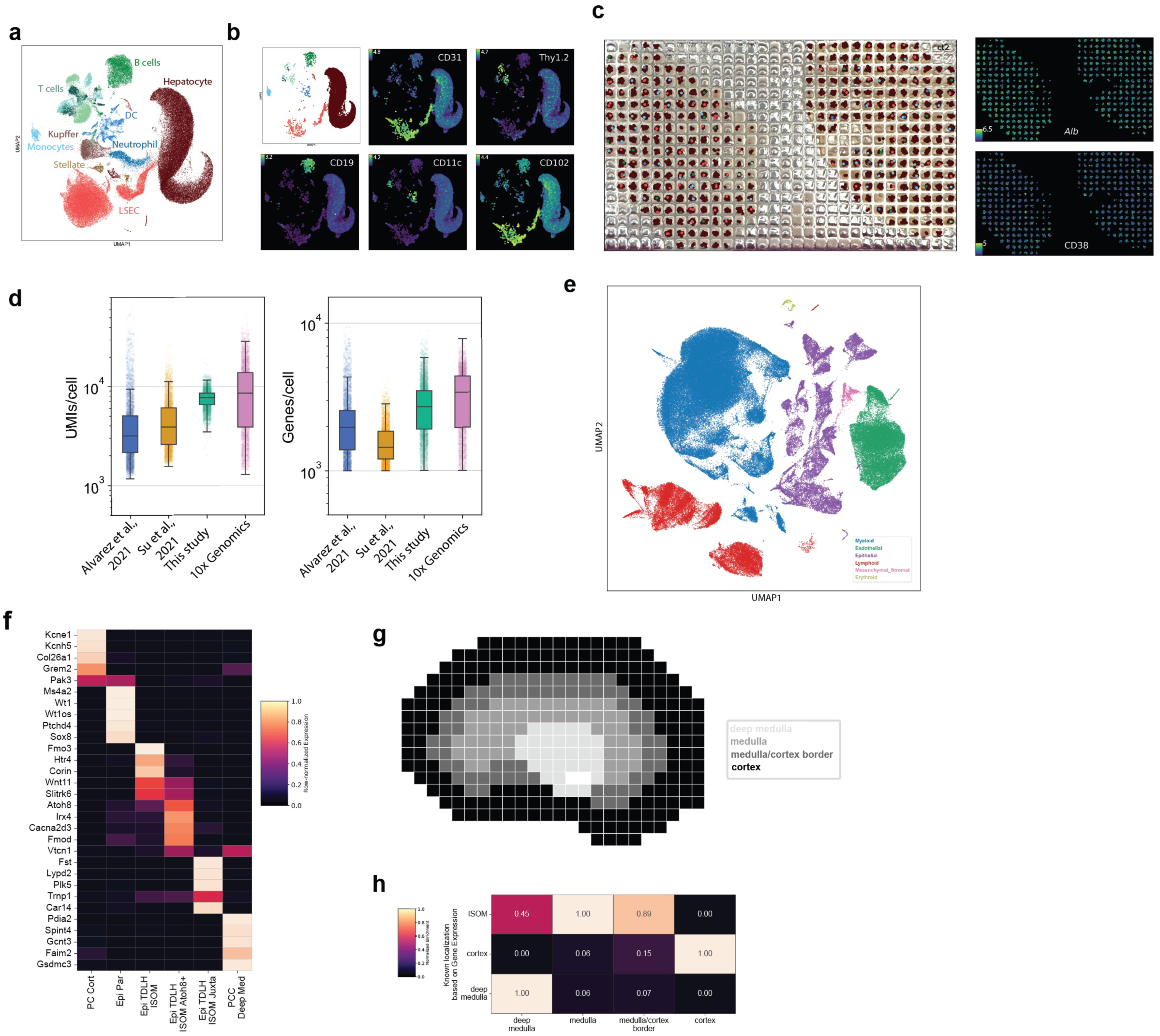
Compatibility of spatial microwell arrays across different tissue types. **a**, Cell type composition of eleven liver sections (115,641 cells) spatially tagged using low-resolution (500µm) microwell arrays. **b**, Lineage markers expressed on the liver main clusters. **c**, Liver spatial maps at 500µm resolution. **d**, Benchmarking of the single-cell transcriptomes obtained from spatially-tagged liver sections against 3 murine liver scRNA-seq datasets^43–45^. **e**, UMAP (based on RNA expression) of 278,261 wild-type kidney cells assayed with 23 arrays with varying resolution. **f**, Cell type-specific gene expression in the kidney^46^. **g**, Manual annotation of kidney spatial regions. **h**, Enrichment of spatially restricted gene expression in defined regions from f. Identity abbreviations are PC, Principal-like Cell; Epi, epithelial; Par, parietal. Location abbreviations are Cort, cortex; TDLH, thin descending loop of Henle; ISOM, inner stripe of the outer medulla; JM, juxtamedullary; DM, deep medulla.

**Extended Data Figure 3.**
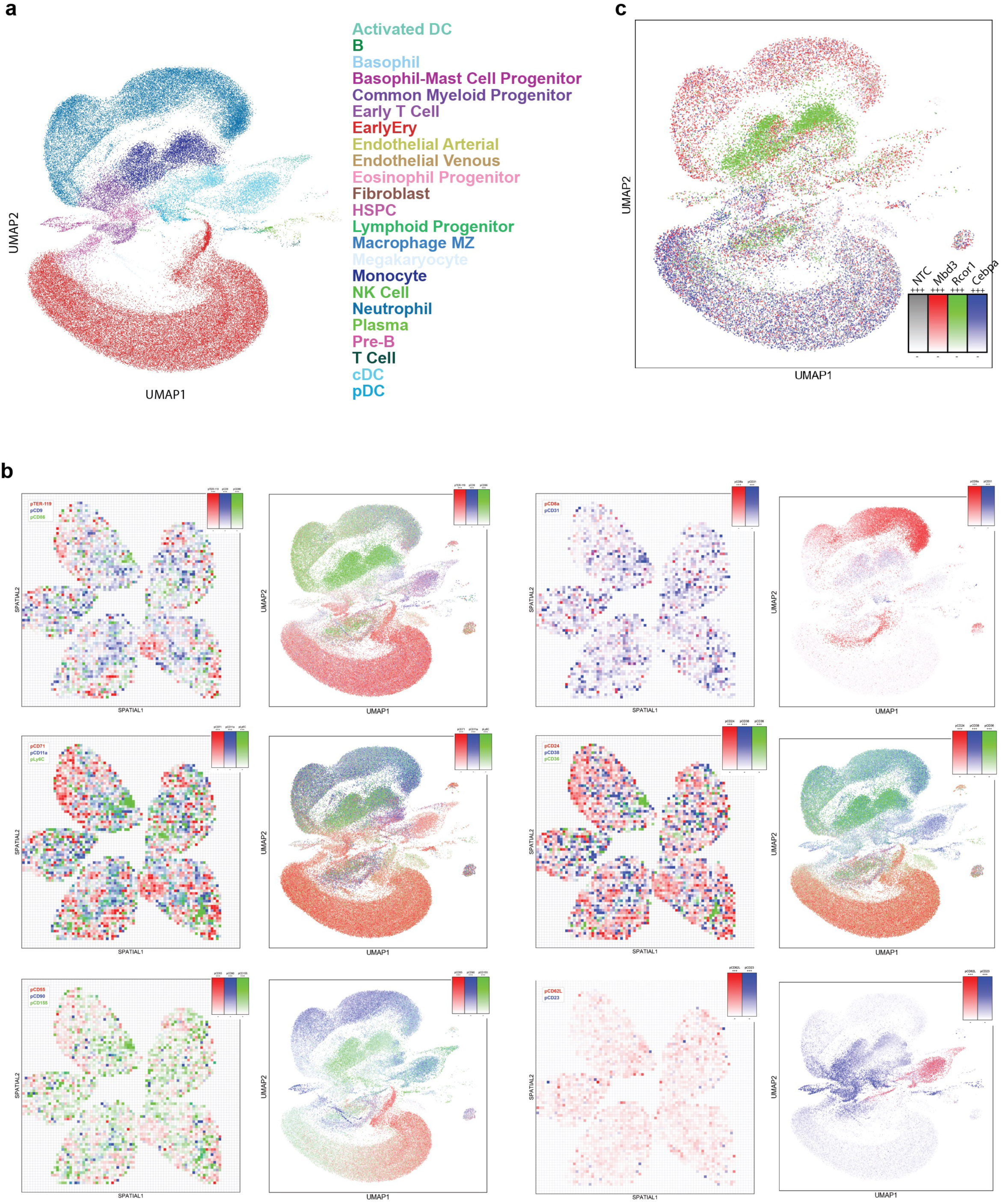
Multimodal PerturbSpace in splenic tissue sections. **a**, UMAP of all profiled splenic cells colored by cell type. **b**, Examples of spatially- and cell type-specific protein expression from CITE-seq library. **c**, UMAP showing differentiation trends for NTC and 3 strong perturbations: *Cebpa*-KO, *Mbd3*-KO and *Rcor1*-KO.

**Extended Data Figure 4.**
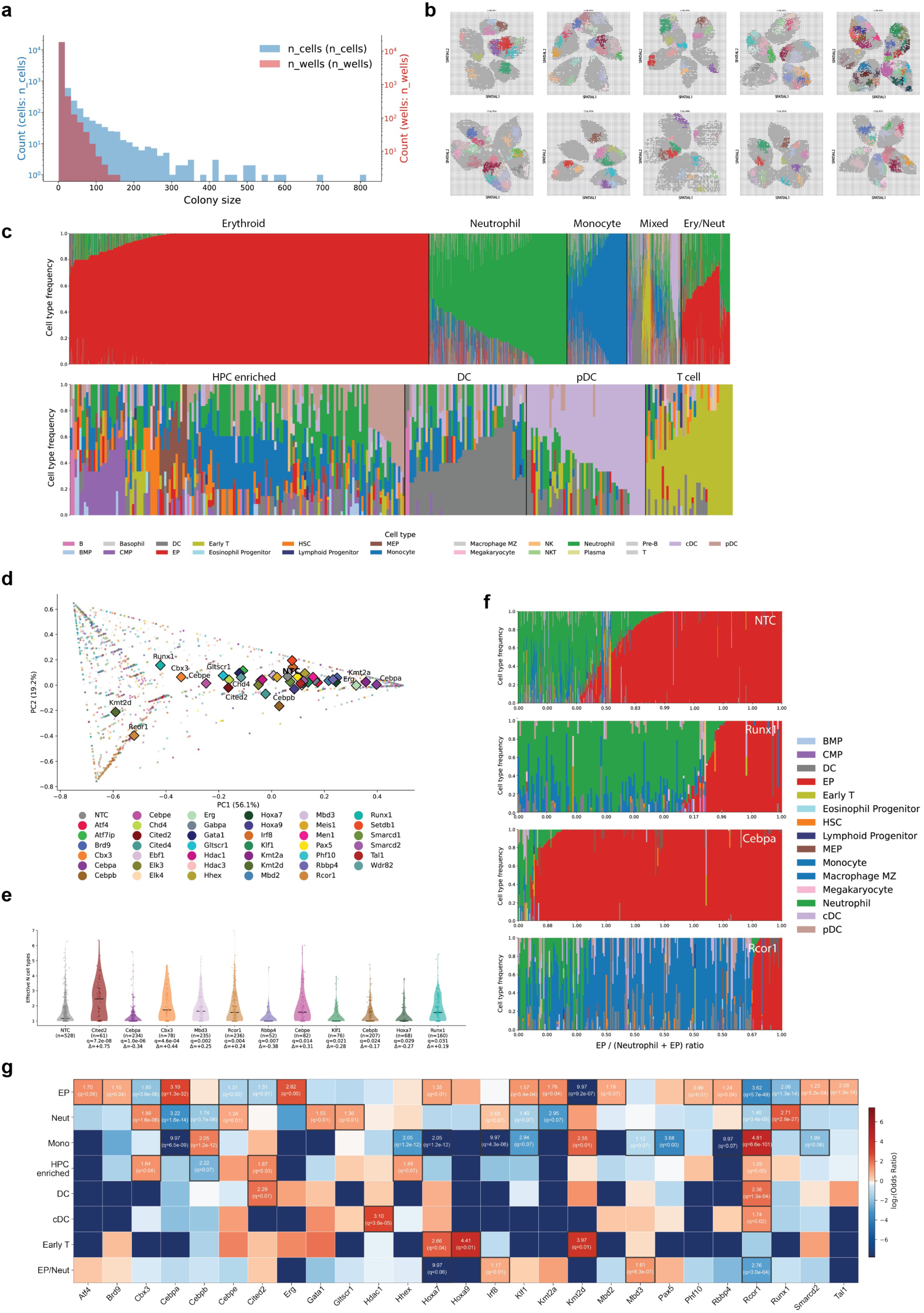
Perturbation-specific effects on splenic colony formation (CFU-S) lineage composition. **a**, Histogram showing the size distribution of CFU-S. Blue bars indicate the total number of cells and red bars indicate the number of spatial neighborhoods (wells) that comprise a CFU-S. **b**, Spleen spatial maps with exemplar CFU-S. **c**, Cell type composition of CFU-S after k-means clustering. Cluster 3 was subclustered revealing four subtypes. **d**, Principal component analysis of lineage colony composition per perturbation. Diamonds represent the centroid for each perturbation. **e**, Perturbation effects on CFU-S diversity. Violin plots showing the number of cell types per colony for each transcriptional perturbation with significantly altered diversity relative to NTC. Certain perturbations such as Cited2-KO increase the lineage diversity in colonies, others such as Cebpa-KO, reduce the diversity shifting composition to uni-lineage colonies. Violin plots show kernel density estimates of the distribution. Black horizontal lines indicate the median. Individual dots represent single colonies. n = number of colonies per perturbation. **f**, Colony lineage compositions for NTC, Runx1-, Cebpa- and Rcor1-KOs. Each bar in the histogram represents an individual colony. Colors reflect colony composition. **g**, Analysis of perturbation effects on colony types highlighting perturbations that enrich or deplete in specific colony types.

**Extended Data Figure 5.**
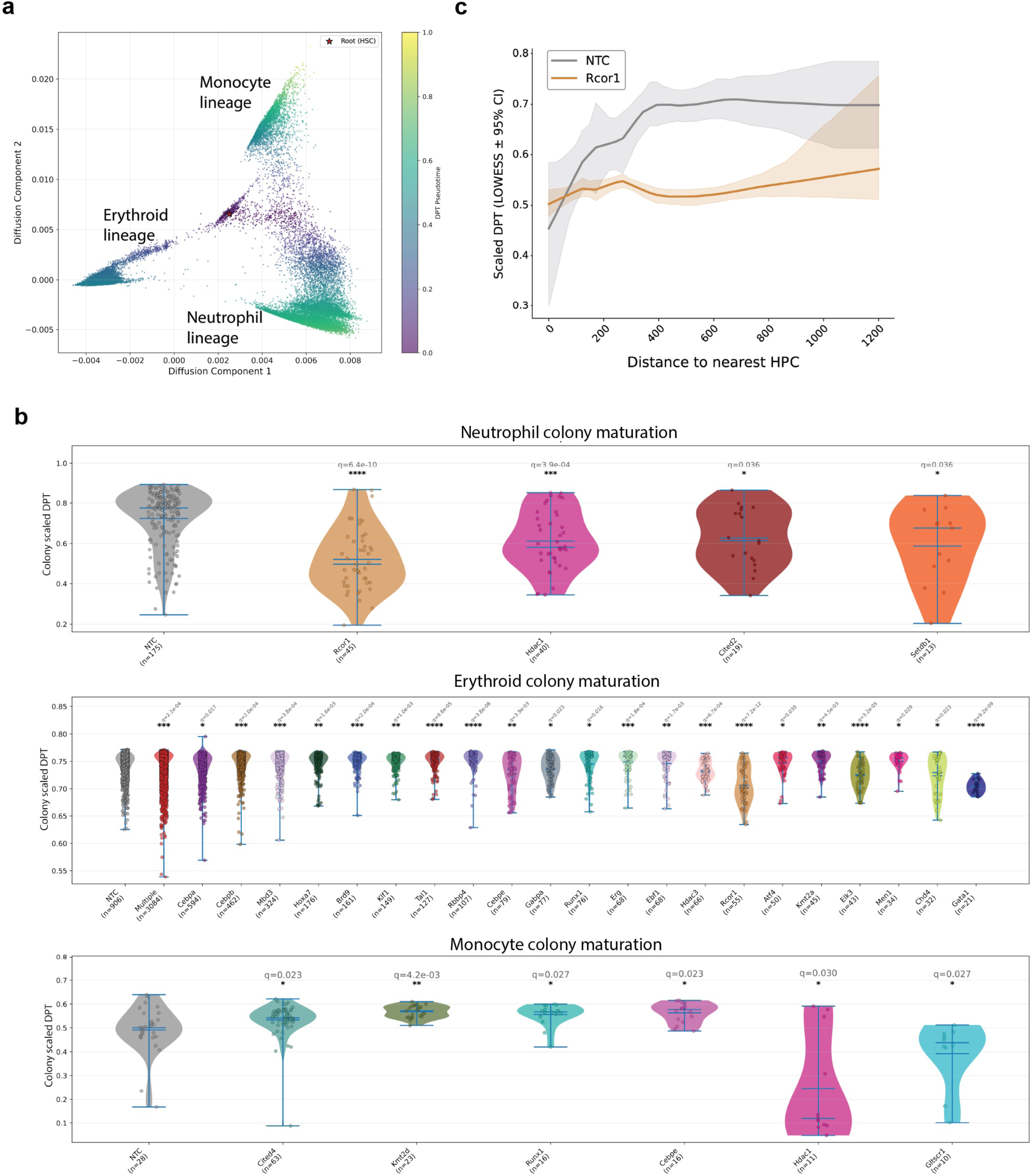
Analysis of colony maturation patterns. **a**, DPT plot displaying the differentiation trajectories in the three most abundant lineages, erythroid, monocyte, neutrophil. **b**, Violin plots showing the effects of transcriptional perturbations on colony maturation patterns (DPT) for neutrophil, monocyte and erythroid types. q ≤ 0.05 vs NTC. Violin plots show kernel density estimates. Center lines indicate median, boxes span the interquartile range, and whiskers extend to 1.5× IQR. Individual dots represent single colonies. **c**, Spatial relationship between the maturation state of colony forming cells and their distance to the founder HPC. Scaled DPT (y-axis) plotted as a function of physical distance to the nearest HPC for Rcor1 knockout (orange) and NTC (gray).

**Extended Data Figure 6.**
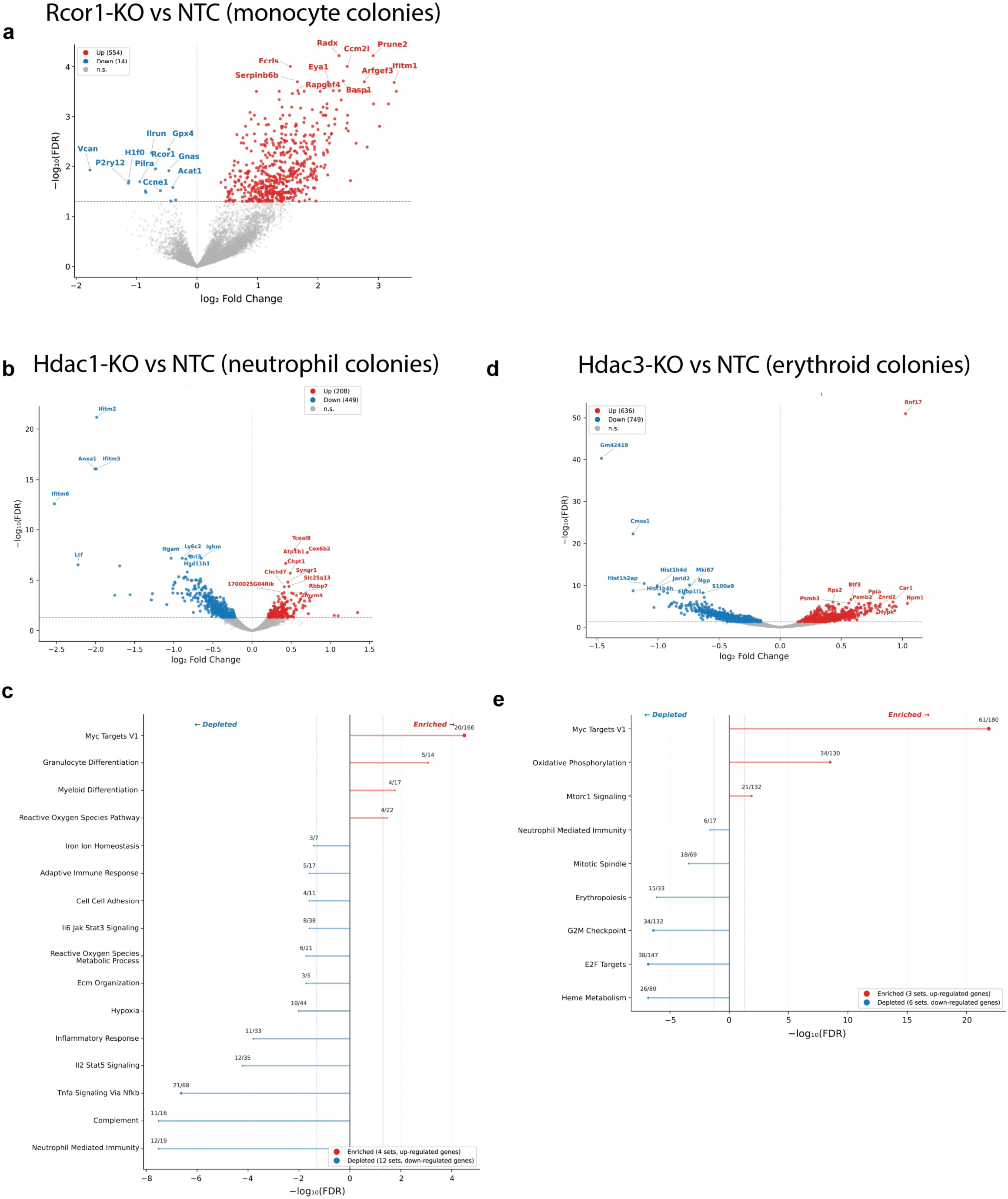
Gene expression analysis in Rcor1-, Hdac1- and Hdac3-perturbed colonies. **a**, Volcano plot showing differential gene expression between Rcor1-perturbed monocyte CFU-S and NTC CFU-S. 554 genes were upregulated and 14 downregulated (FDR < 0.05). ORA plot is presented in Fig. 2g. **b**, Volcano plot for Hdac1 perturbed neutrophil colonies. 208 genes were upregulated and 449 genes were downregulated compared to NTC. **c**, ORA plot for differentially expressed genes in b. FDR < 0.05. **d**, Volcano plot for Hdac3 perturbed erythroid progenitor colonies. 636 genes were upregulated and 749 genes were downregulated compared to NTC. **e**, ORA plot for differentially expressed genes in d. FDR < 0.05.

**Extended Data Figure 7.**
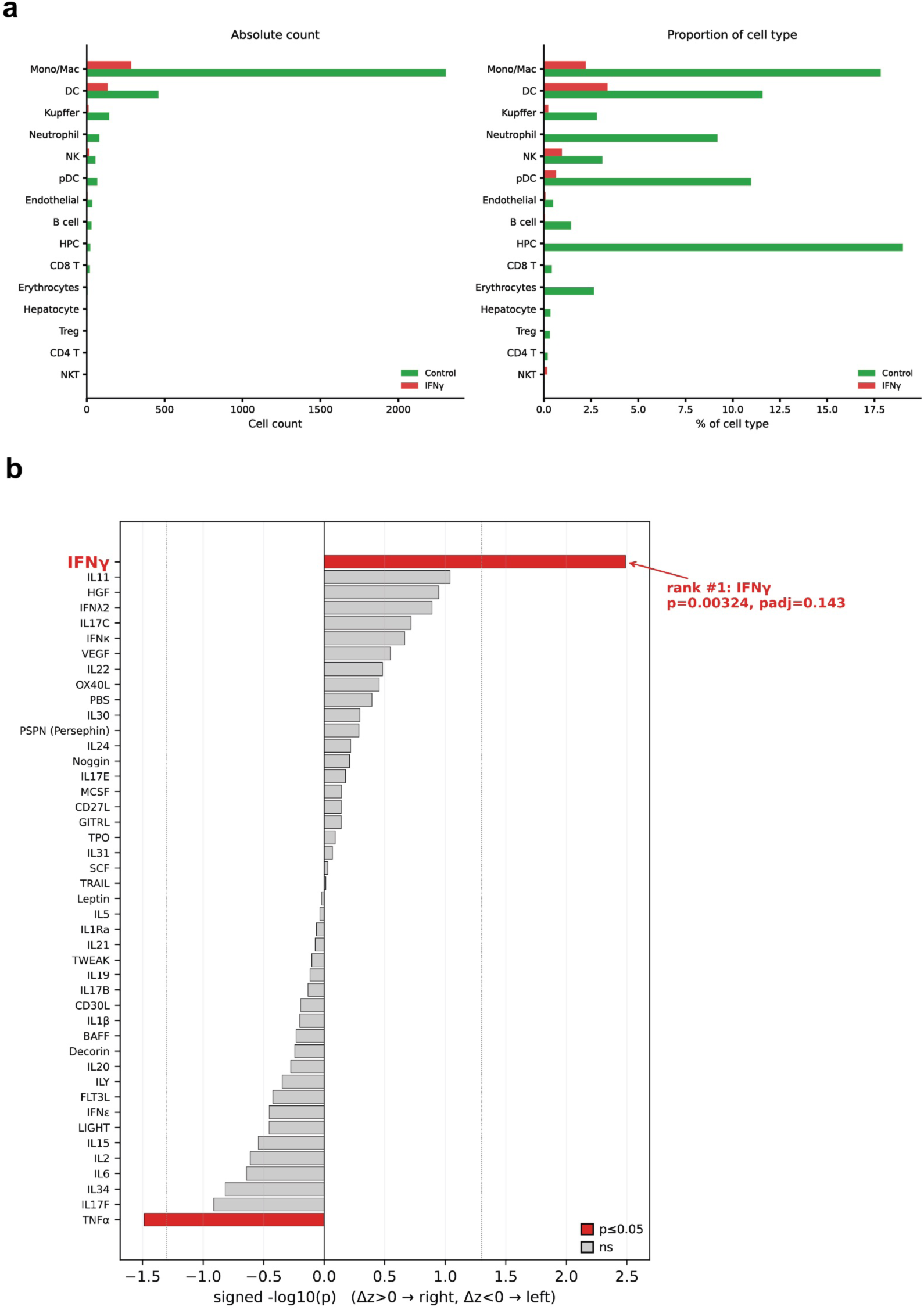
Analysis of non-cell autonomous effects driven by IFNγ- expressing cells in the liver. **a**, Number and proportion of IFNγ or Ctrl expressing cells per cell type. **b**, Full cytokine activity response programs from Fig. 3.

**Supplemental Figure 1.**
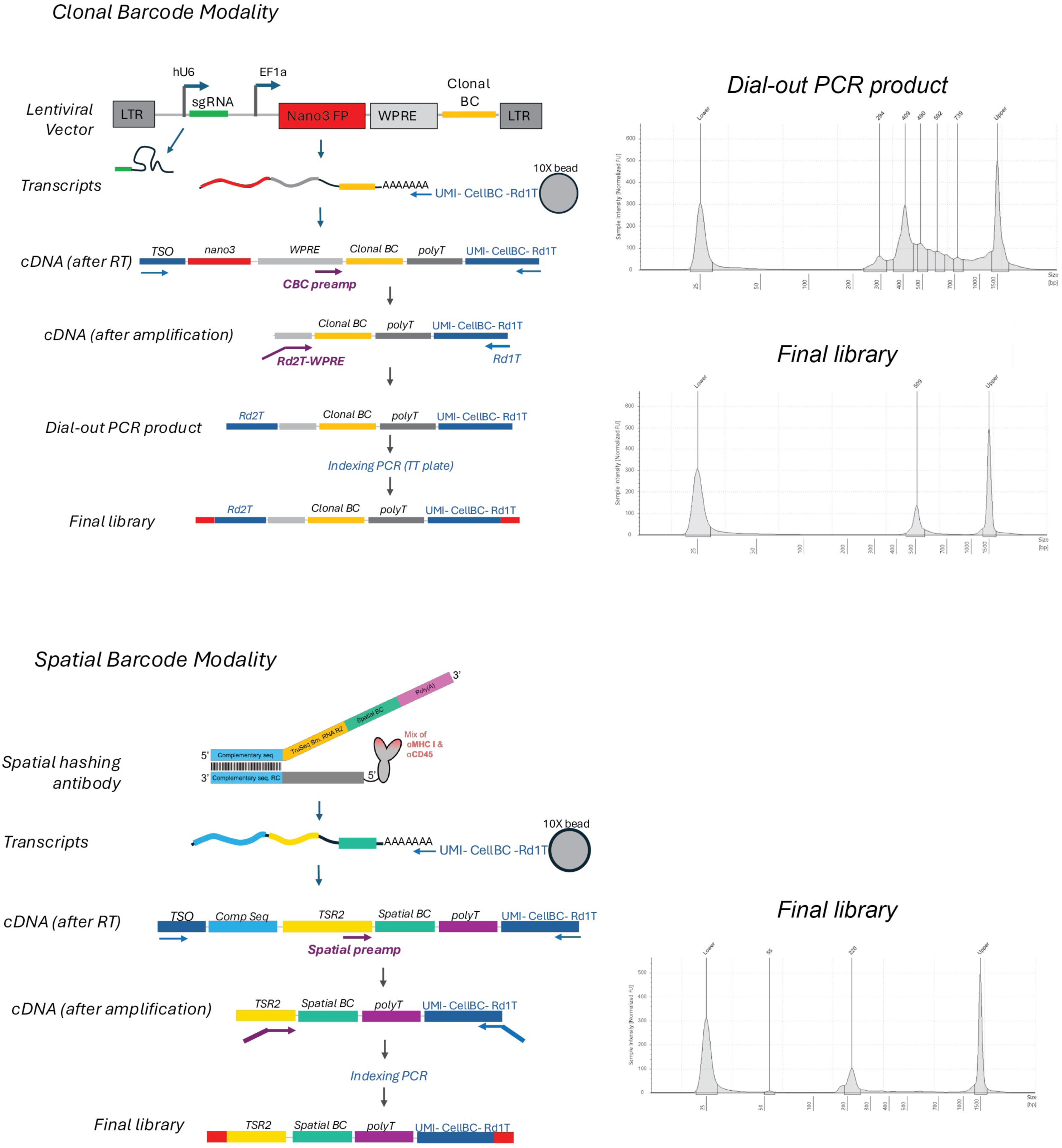

